# Shift of maternal gut microbiome of Tibetan antelope (*Pantholops hodgsonii*) during the perinatal period

**DOI:** 10.1101/2020.01.13.903591

**Authors:** Yue Shi, Ziyan Miao, Jianping Su, Samuel K. Wasser

**Affiliations:** Department of Biology, University of Washington, Box 351800, Seattle, WA 98195, USA; Qinghai Key Laboratory of Animal Ecological Genomics, Northwest Institute of Plateau Biology, Chinese Academy of Sciences, No. 23 Xinning Road, Xining, Qinghai 810008, China; Museum of Natural Resources of Qinghai Province, Xining, Qinghai 810008, China

**Author notes:** Correspondence: Yue Shi. Wisconsin Cooperative Fishery Research Unit, University of Wisconsin - Stevens Point, Stevens Point, WI 54481. Deceased on 27 June 2018 (Jianping Su).

**Keywords:** maternal gut microbiome, perinatal period, hormone, Tibetan antelope, reproductive health

## Abstract

The maternal gut microbiome can influence and be affected by the substantial physiological changes taking place during the perinatal period. However, little information is known about the changes in the maternal gut microbiome during this period. Tibetan antelope (*Pantholops hodgsonii*) provide a unique system to address this issue because their summer migration cycle is synchronized with the perinatal period. We used 16S rRNA gene sequencing to generate gut microbiome profiles using fecal samples collected from female migratory Tibetan antelope. We then correlated microbiome diversity with fecal hormone metabolite concentrations of glucocorticoids (GCs) and triiodothyronine (T3) extracted from the same fecal samples. The maternal gut microbiome of Tibetan antelope was dominated by Firmicutes and Bacteroidetes. There was a clear separation in gut microbial composition by female reproductive states based on both hierarchical clustering and PCoA analyses. The shift in the maternal gut microbiome likely reflects the metabolic and immune system dynamics during the perinatal period. Overall, the microbiome diversity was higher in the late pregnancy compared to the postpartum period. The negative association between T3 and microbiome diversity may be moderated by the shift of reproductive states since the correlations disappeared when considering each reproductive state separately. Integrating the microbiome dimension, migration pattern and reproduction may have direct conservation implications as by establishing a baseline of the physiological changes during the migration/perinatal period, we can have a better understanding of the impacts of increasing human activities on the Tibetan Plateau on the reproductive health of Tibetan antelope.

## Introduction

The transition from pregnancy to lactation during the perinatal period is among the most important determinants of maternal-offspring health outcomes (Blaser & Domínguez-Bello, 2016; Dunlop et al., 2015; Prince et al., 2015). Substantial physiological changes occur during this transition, including changes in hormones, immune system and metabolism, to shift resource allocation from energy storage to milk synthesis and preserve the health of both the mother and her offspring (Lain & Catalano, 2007; Zeng, Liu, & Li, 2017). Both late pregnancy and lactation are the two most energetic demanding female reproductive periods, with even more pronounced energy demands during lactation (Butte & King, 2005). Hormones serve as mediators in the process, directing nutrients and energy to the highly specialized maternal reproductive tissues and the developing fetus (Picciano, 2003). Two important metabolic hormones are involved in this transition, including glucocorticoids (GCs) and thyroid hormones (THs). GCs are released from the adrenal glands in vertebrates and regulated by the hypothalamic-pituitary-adrenal (HPA) axis. GCs can rapidly mobilize glucose in response to physiological and psychological stress (Palme, Rettenbacher, Touma, EL-Bahr, & Mostl, 2006). GC levels are elevated during pregnancy, allowing for greater substrate availability for fetal growth (Lain & Catalano, 2007; Zeng et al., 2017). THs are released by the thyroid gland and regulated by the hypothalamus-pituitary-thyroid gland axis. THs function as a metabolic thermostat, continuously monitoring energy intake and regulating metabolism accordingly (Douyon & Schteingart, 2002). Thyroid activity increases throughout the pregnancy. During the transition from pregnancy to lactation, metabolic adjustments of thyroid hormones are essential in establishing metabolic priority for the lactating mammary gland (Capuco, Connor, & Wood, 2008).

The maternal gut microbiome is likely to influence and be affected by the physiological changes during the perinatal period and it is critical for the establishment and development of the neonatal microbiome (Prince et al., 2015). Pregnancy is associated with a profound alteration of maternal gut microbiota (Koren et al., 2012). However, little research has been conducted to assess how maternal gut microbiota changes during the transitionary perinatal period. Two such studies focused on human microbiome but had inconsistent results: one study found that human maternal gut microbiota remained stable over the perinatal period (Jost, Lacroix, Braegger, & Chassard, 2013) and the other study concluded that there was a change in microbial community structure and reduced microbiome diversity from pregnancy to postpartum in humans (Crusell et al., 2018). A third study, which was conducted on dairy cows, revealed distinct microbiome profiles between pre- and postpartum females and argued that these changes resulted from the shifts in diet regime (Lima et al., 2015). More information is needed to document the shift in the microbiota composition during this critical transition period, which has profound impacts on the health outcomes of both females and offspring.

Gut microbiome studies have been conducted predominantly on humans and nonhuman primates (West et al., 2019). Comprehensive surveys of the microbiome composition in non-model species remain relatively rare, with primary focuses on the species of economic importance, such as cattle and horse (O’ Donnell, Harris, Ross, & O’Toole, 2017). Microbiome research in wildlife tend to focus on the impacts of land-use change, climate change, captive breeding, invasive species, antibiotics, infectious disease and environmental contamination on the microbiome diversity and community structure (Redford, Segre, Salafsky, del Rio, & McAloose, 2012; Trevelline, Fontaine, Hartup, & Kohl, 2019). We found no research integrating the microbiome dimension into wildlife reproduction.

Tibetan antelope (*Pantholops hodgsonii*) provide a unique system to study changes in the microbiome during the perinatal period. Every summer, female Tibetan antelope depart from the wintering sites to the calving sites in May - June and return with their newborns in late July - early August (Schaller, 1998). Their summer migration cycle is synchronized with the perinatal period (Buho et al., 2011) with considerable energy demand when both the females and offspring are most vulnerable (Butte & King, 2005). Tibetan antelope were reduced to the brink of extinction at the end of the 20^th^ century by illegal poaching for their underfur. International conservation efforts successfully curbed the poaching through law enforcement and habitat protection. Their population size has recovered since 2011 (Leclerc, Bellard, Luque, & Courchamp, 2015), but recent habitat fragmentation, such as fencing and road construction, could impede the recovery of Tibetan antelope. Integrating the microbiome dimension, migration pattern and reproduction may have direct conservation implications as by establishing a baseline of the physiological changes during the migration/perinatal period, we can have a better understanding of the impacts of increasing human activities on the Tibetan Plateau on the reproductive health of Tibetan antelope.

The aim of this study was to characterize the changes in the maternal gut microbiome of Tibetan antelope during the perinatal period by using 16S rRNA gene sequencing technology and explore the relationships between the changes in the gut microbiome diversity and metabolic hormones, GC and triiodothyronine (T3), the most biologically active form of THs. We used fecal samples as a proxy for the gut microbiome due to their accessibility and non-invasive nature (Yasuda et al., 2015), though fecal samples are not completely representative of the entire gut microbiome (Ingala et al., 2018). The same fecal samples were also used to extract fecal hormone metabolites. We predicted that 1) the maternal gut microbiome of Tibetan antelope is dominated by phyla Firmicutes and Bacteroidetes as with other mammals (Ley et al., 2008); 2) there is a shift in microbiome community composition in the perinatal period in response to the metabolic adjustments during the same period. To control for the potential confounding impact of dietary changes during the sampling period on the maternal gut microbiome composition, we included the sampling time as a factor when assessing the changes in microbial community composition. Our study sheds light on how the microbiome composition shifts during this transition period and emphasizes the importance of incorporating the maternal gut microbiome into the conservation management efforts to support animal reproductive health and overall recovery success of Tibetan antelope.

## Materials and Methods

### Ethics Statement

Tibetan antelope is listed in the Category I of the National Key Protected Wild Animal Species under China’s Wild Animal Protection Law. In September 2016, Tibetan antelope were reclassified from Endangered to Near Threatened by the International Union for Conservation of Nature (IUCN) Red List due to the recovery of their population size. Sample collection and field studies adhered to the Wild Animals Protection Law of the People’s Republic of China. Fresh scat samples were collected under IACUC protocol #2850-12 and local regulations to minimize disturbance.

### Sample Collection

Qinghai-Tibet Railway bisects the migration route of Tibetan antelope approximately 40 km from their summer calving area at Zhuonai Lake. Female Tibetan antelope almost exclusively use the Wubei Bridge underpass (35°15’2.71”N, 93° 9’45.12”E), a 198m long, 30m wide structure (Xia, Yang, Li, Wu, & Feng, 2007). We collected fecal samples at the Wubei Bridge underpass when females migrated to and returned from the calving ground in 2017. Females are in the late pregnancy stage when migrating to the calving ground, and in the postpartum period when on their return to the wintering ground. Animals were observed with binoculars from a recommended viewing distance of ∼ 300 m (Lian, Zhang, Cao, Su, & Thirgood, 2007) until they defecated and left the area. To minimize the chance of collecting multiple fecal samples from the same individual, three field assistants were involved in sample collection in three different directions. We excluded fecal samples of small size to avoid collecting fecal samples from the young and newborns. Adult males do not conduct seasonal migration. A total of 65 fresh scat samples were collected and placed into individual zip-lock bags along with records of the date and GPS coordinates. Fifty samples were collected during the late pregnancy stage and 15 samples collected during the postpartum period. Samples were kept frozen at – 20 °C until lab analyses.

### DNA Extraction and Species Identification

To minimize environmental contamination, we extracted DNA from the well-preserved core of one fecal pellet per sample (Menke, Meier, & Sommer, 2015) using the E.Z.N.A.^®^ DNA isolation kit (Omega Biotek, Norcross, GA, U.S.). The DNA concentration and quality were evaluated using Nanodrop™ 2000 Spectrophotometer (Nanodrop, Wilmington, DE, USA), and visualized with 1% agarose gel electrophoresis. A short fragment (196 bp) of mtDNA cytochrome c oxidase subunit I gene was amplified and sequenced using the forward primer (5’ GCCCCTGATATAGCATTCCC 3’) and the reverse primer (5’ CTGCCAGGTGTAGGGAGAAG 3’). PCR was conducted using EasyTaq PCR SuperMix (Transgen Biotech, Inc.) with 4 ul DNA template. 2.5 ul 10 mg/ml of bovine serum albumin was added to the PCR mix to improve amplification success. We followed the recommended thermo protocol in the kit with an annealing temperature of 51° C. Sequences were checked against the NCBI database using BLAST for species confirmation.

### Library Preparation

The V3-V4 region of the 16S rRNA gene was amplified using the forward primer 338F (5’-ACTCCTACGGGAGGCAGCAG-3’) and the reverse primer 806R (5’-GGACTACHVGGGTWTCTAAT-3’) (Mori et al., 2014). Each primer was designed to contain: 1) the appropriate Illumina adapter sequence allowing amplicons to bind to the flow cell; 2) an 8bp index sequence; and 3) gene-specific primer sequences as described above. PCR reactions were conducted using the TransStart^®^ FastPfu DNA Polymerase kit (TransBionova Co., Ltd, Beijing, China) with 20 μL reaction solution in total including 10 ng template DNA, 4 μL 5×FastPfu buffer, 2 μL 2.5 mM dNTPs, 0.8 μL of each primer (5 μM), 0.4 μL FastPfu Polymerase and 0.2 μL 20 ng/ μL of bovine serum albumin. The PCR cycling conditions were as follows: 95 °C for 3 min, followed by 27 cycles of 95 °C for 30 s, 55 °C for 30 s, and 72 °C for 45 s and a final extension of 72 °C for 10 min. All PCR products were visualized on agarose gels (2% in TAE buffer) and purified with AxyPrep DNA Gel Extraction Kit (Axygen Biosciences, Union City, CA, USA). Before sequencing, DNA samples were quantified using QuantiFluor™-ST (Promega, USA). Paired-end amplicon libraries were constructed, and sequencing was performed using the Illumina MiSeq PE300 platform at Majorbio BioPharm Technology Co., Ltd., Shanghai, China.

### Bioinformatics Analyses

Sequencing reads were analyzed using the *DADA2* pipeline (Callahan et al., 2016) with the following steps: 1) Initial quality filtering was performed using the *filterAndTrim* command with default parameters. Forward and reverse reads were truncated at 290 bp and 210 bp respectively; 2) Amplicon sequence variants (ASVs) were inferred using the core DADA2 sample inference algorithm, *dada*, after error rate model generation and sequence dereplication; 3) Forward and reverse reads were merged to obtain the full denoised sequences using the *mergePairs* command; 4) Chimeric sequences were identified and removed using the *removeBimeraDenovo* command; 5) Taxonomy classification was assigned to each ASV using the *assignTaxonomy* command using the Silva reference database (version 132/16s_bacteria) (Quast et al., 2012). 16S rRNA gene analysis is limited for species-level classification because closely related bacterial species may have near-identical 16S rRNA gene sequences (Plummer & Twin, 2015). Therefore, each ASV was assigned only down to the genus level. Finally, ASVs with less than 10 copies in the resulting ASV table were removed.

### Fecal Hormone Metabolite Analyses

Fecal samples were freeze-dried to remove water content. Fecal GC and T3 metabolites were then extracted from freeze-dried fecal samples following previously described protocols (Wasser et al., 2010; 2004; 2000). Briefly, each freeze-dried fecal sample was thoroughly homogenized before suspending ∼0.1 g of fecal powder in 15 ml of 70% ethanol, followed by 30 min continuous agitation. After centrifugation (1800 g; 20 min), the supernatant was poured into a new vessel. An additional 15 ml of 70% ethanol was added to the remaining fecal power for a second extraction. This mixture was agitated for 30 min and centrifuged at 1800 g for 20 min. Supernatants from both extractions were combined and stored at −20 °C.

Since most native hormones are excreted as metabolites in feces (Palme, Fischer, Schildorfer, & Ismail, 1996), it is essential to select an appropriate assay system that includes a group-specific antibody, which is capable of detecting the predominant fecal metabolites of the parent hormone in the species investigated (Palme et al., 1996; Touma & Palme, 2006; Wasser et al., 2000). Fecal GC metabolites were diluted with assay buffer at the ratio of 1:30 and analyzed using DetectX^®^ Corticosterone Immunoassay kit (Arbor Assays). This assay kit was chosen because a previous study has shown that corticosterone antibody has the highest affinity for the major cortisol metabolites (Wasser et al., 2000). Fecal T3 metabolites were diluted at the ratio of 1:60 and analyzed using DetectX^®^ Triiodothyronine (T3) Immunoassay kit (Arbor Assays) because T3 metabolite maintains its pure form in many mammalian species (Wasser et al., 2010). Both hormone assays were validated using serially diluted fecal extracts pooled from 10 random samples. The displacement curves were parallel to those of standard hormone preparations, indicating that hormone concentrations were reliably measured across their ranges of concentrations. Besides, the recovery rate of the GC and T3 standards was 127.8% and 127.6% respectively, indicating a limited amount of interference in hormone binding from the substances in the fecal extracts of Tibetan antelope.

### Statistical Analyses

Rarefaction curves were plotted for each sample using the *rarecurve* function in the *vegan* R package to assess the adequacy of sequencing depth. Variations in sequencing depth among samples were normalized using the variance stabilizing transformation (VST) implemented in the *DESeq2* R package (McMurdie & Holmes, 2014). Hierarchical clustering and principal coordinate analyses (PCoA) based on the Euclidean distance matrix of VST data were performed to visualize microbial composition differences as a function of reproductive states (late pregnancy vs. postpartum period). Analysis of Similarity (ANOSIM) test was used to assess significant differences in microbial community composition as a function of reproductive states while controlling for the time of the year using *strata*. We filtered out phyla, classes, and orders with relative abundance less than 0.1% and familiae and genera with a relative abundance of less than 1%. Changes in the relative abundance of taxa between reproductive states were analyzed through Wilcoxon signed-rank tests and *p* values were adjusted with the Benjamini-Hochberg method to control for false discovery rate. We used the *Phyloseq* R package to compute alpha diversity including richness measurements (Observed number of ASVs, Chao 1 richness estimator, Abundance-Based Coverage Estimator or ACE) and diversity indices (Shannon diversity index, Inverse of the Simpson diversity estimator or InvSimpson, Fisher’s index). T-tests were conducted to compare microbial alpha diversity between reproductive states. Fecal hormone metabolite concentrations were natural log-transformed to meet the assumption of normality. We used Wilcoxon signed-rank tests to compare fecal hormone metabolite concentrations between reproductive states since the assumption of homogeneity of variance was not met. We constructed linear models of microbial alpha diversity as a function of fecal hormone metabolite concentrations (natural log-transformed), reproductive states and interactions between main effects. We used analysis of covariance (ANCOVA) to compute the analysis of covariance tables for fitted models.

## Results

### Maternal gut microbiota is dominated by Firmicutes and Bacteroidetes

After quality filtering, denoising, read merging and singleton/chimera removal, 1,415,743 high-quality sequence reads were retained with an average of 21,781 reads per sample (from 15,129 to 32,148) and a median sequence length of 406 bp. In total, there were 6,026 ASVs identified. The rarefaction curves on the number of ASVs reached a plateau, suggesting that the sequencing depth was adequate, and the microbial community was well surveyed (Supplementary Figure 1). Taxonomic assignments revealed 8 bacterial phyla with relative abundance above 0.1% (Figure 1). Dominant phyla were Firmicutes (67.97%), Bacteroidetes (26.30%), Verrucomicrobia (2.94%) and Actinobacteria (1.12%). There were 11 classes with relative abundance above 0.1% with dominant classes including Clostridia (phylum: Firmicutes; 66.92%), Bacteroidia (phylum: Bacteroidetes; 26.30%), Verrucomicrobiae (phylum: Verrucomicrobia; 2.94%) and Actinobacteria (phylum: Actinobacteria; 1.03%) (Supplementary Figure 2). All sequence reads were assigned at the levels of phylum and class, suggesting there were few sequencing artifacts in the dataset. At the order level, we found 11 orders with the relative abundance above 0.1%, with dominant orders including Clostridiales (phylum: Firmicutes; 66.89%), Bacteroidales (phylum: Bacteroidetes; 26.06%), Verrucomicrobiales (phylum: Verrucomicrobia; 2.93%) and Micrococcales (phylum: Actinobacteria; 1.03%) (Supplementary Figure 3). Ten familiae with relative abundance greater than 1% were found (Figure 2). Dominant familiae included *Ruminococcaceae* (phylum: Firmicutes; 46.79%), *Lachnospiraceae* (phylum: Firmicutes; 12.17%), *Bacteroidaceae* (phylum: Bacteroidetes; 9.21%), *Rikenellaceae* (phylum: Bacteroidetes; 8.42%) and *Christensenellaceae* (phylum: Firmicutes; 5.79%), *Akkermansiaceae* (phylum: Verrucomicrobia; 2.93%), *Prevotellaceae* (phylum: Bacteroidetes; 2.80%), *Muribaculaceae* (phylum: Bacteroidetes; 1.21%) and *Micrococcaceae* (phylum: Actinobacteria; 1.03%). About 4.41% of sequence reads could not be assigned at the family level. The taxonomic classification at the genus level is ambiguous as about 21.58% of sequence reads could not be assigned at the genus level (Supplementary Figure 4).

**Figure 1.**
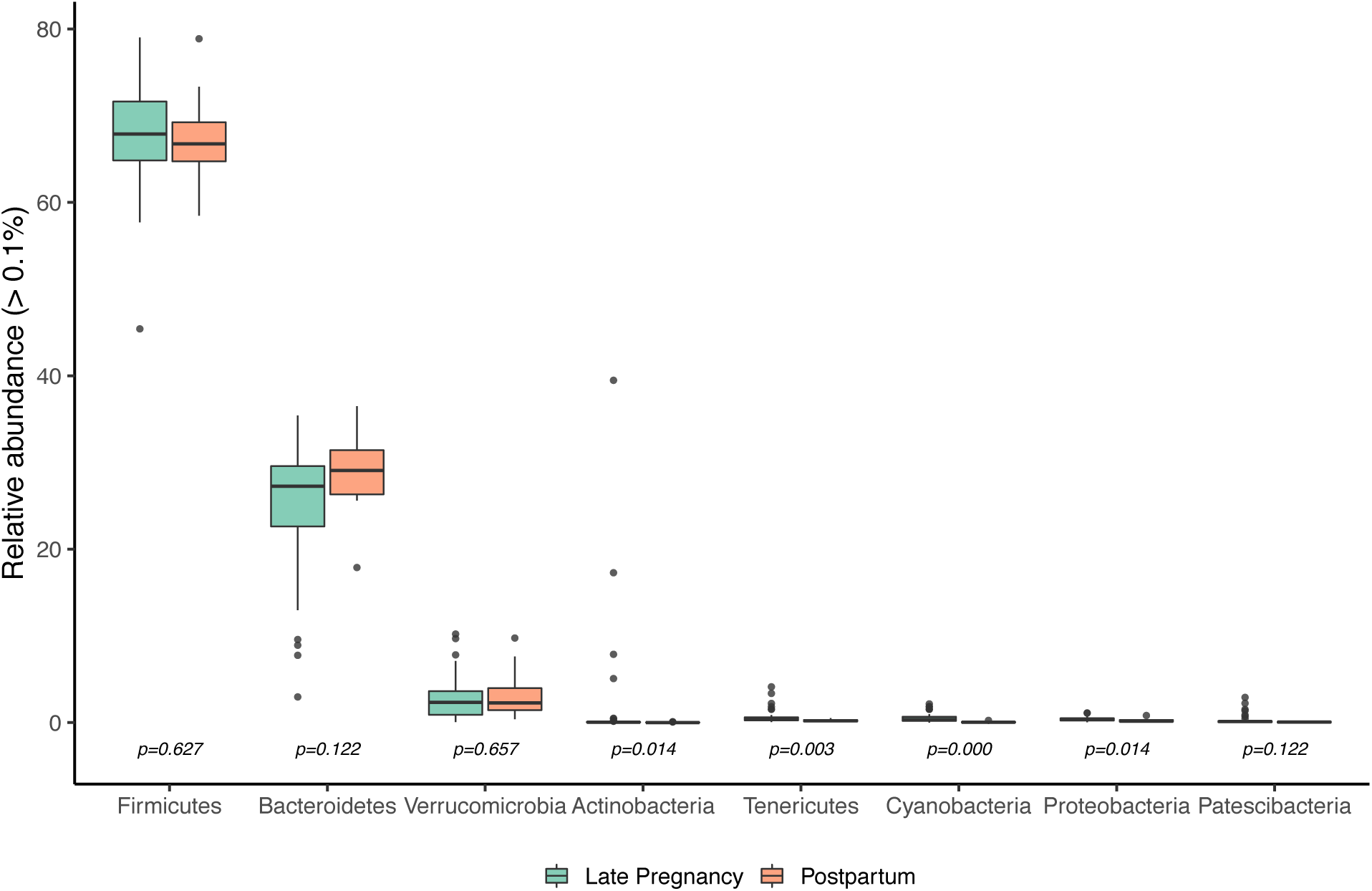
Phyla found in the maternal gut microbiome of Tibetan antelope with relative abundance greater than 0.1%. Changes in the relative abundance of phyla between reproductive states were analyzed through Wilcoxon signed-rank tests and p values were adjusted with the Benjamini-Hochberg method to control for false discovery rate.

**Figure 2.**
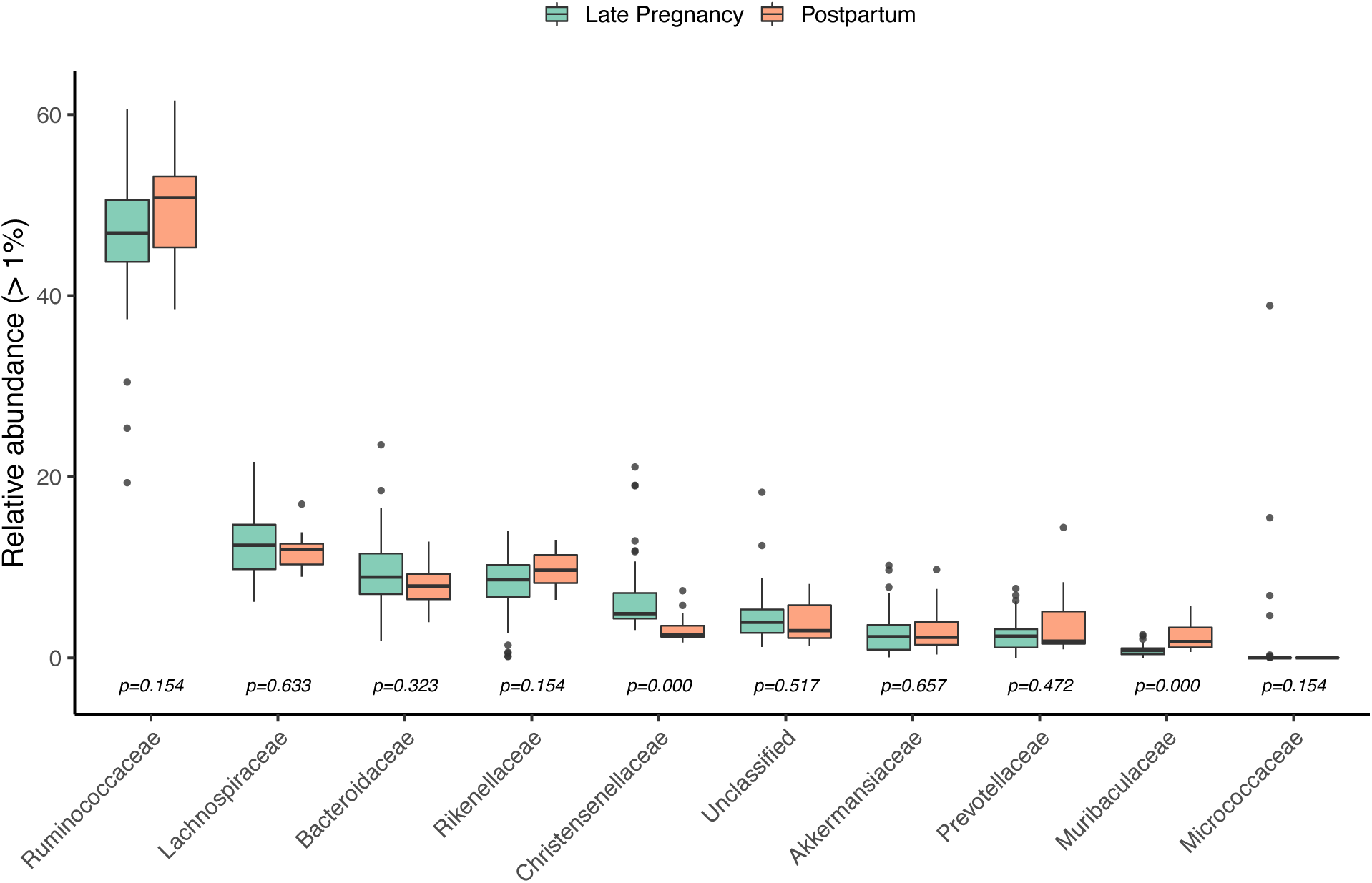
Familiae found in the maternal gut microbiome of Tibetan antelope with relative abundance greater than 1%. Changes in the relative abundance of familiae between reproductive states were analyzed through Wilcoxon signed-rank tests and p values were adjusted with the Benjamini-Hochberg method to control for false discovery rate.

**Figure 3.**
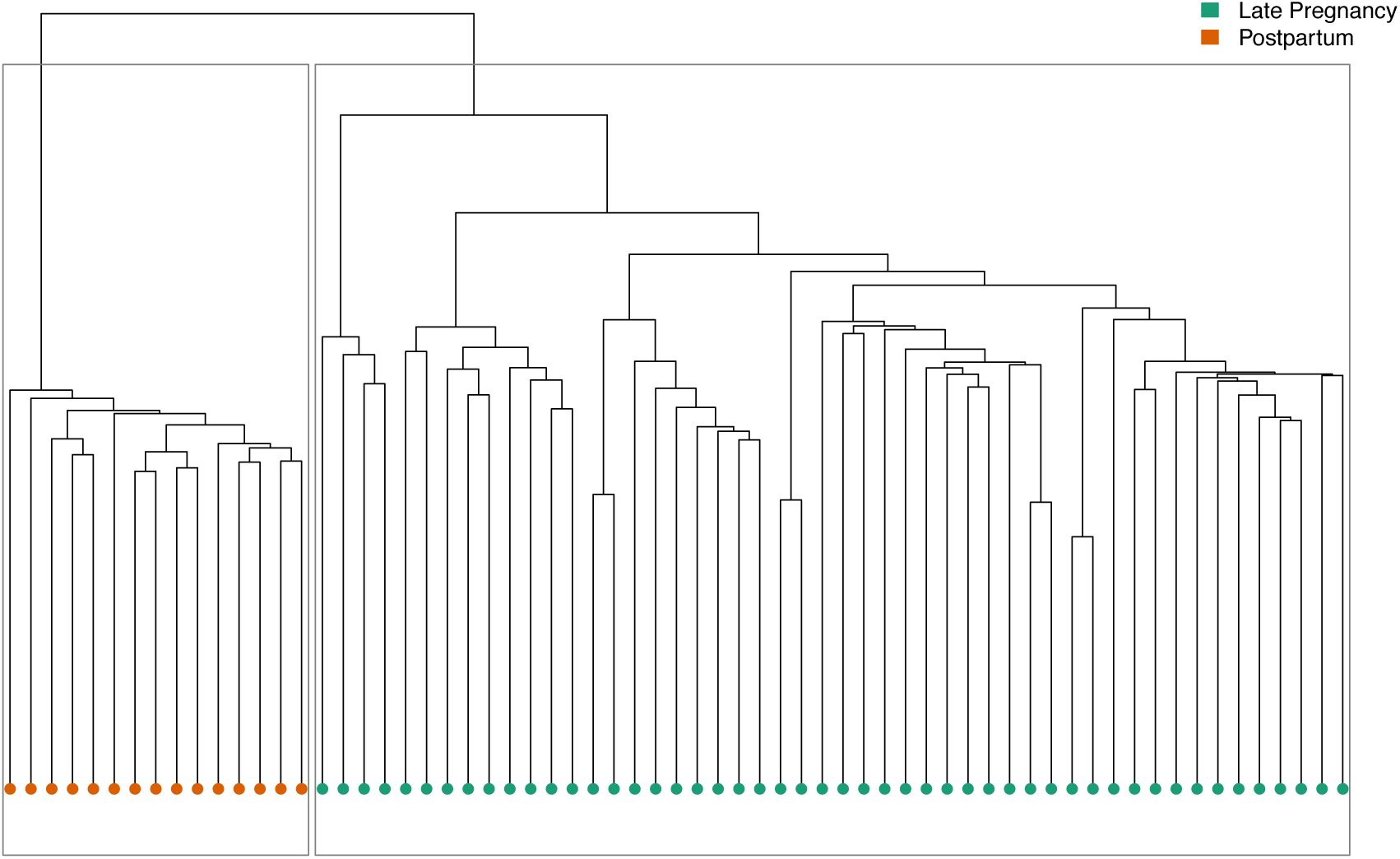
Hierarchical clustering analysis of maternal gut microbiota in female Tibetan antelope with a Euclidean distance matrix after variance stabilizing transformation. Each leaf of the dendrogram represents one fecal sample (N=65).

**Figure 4.**
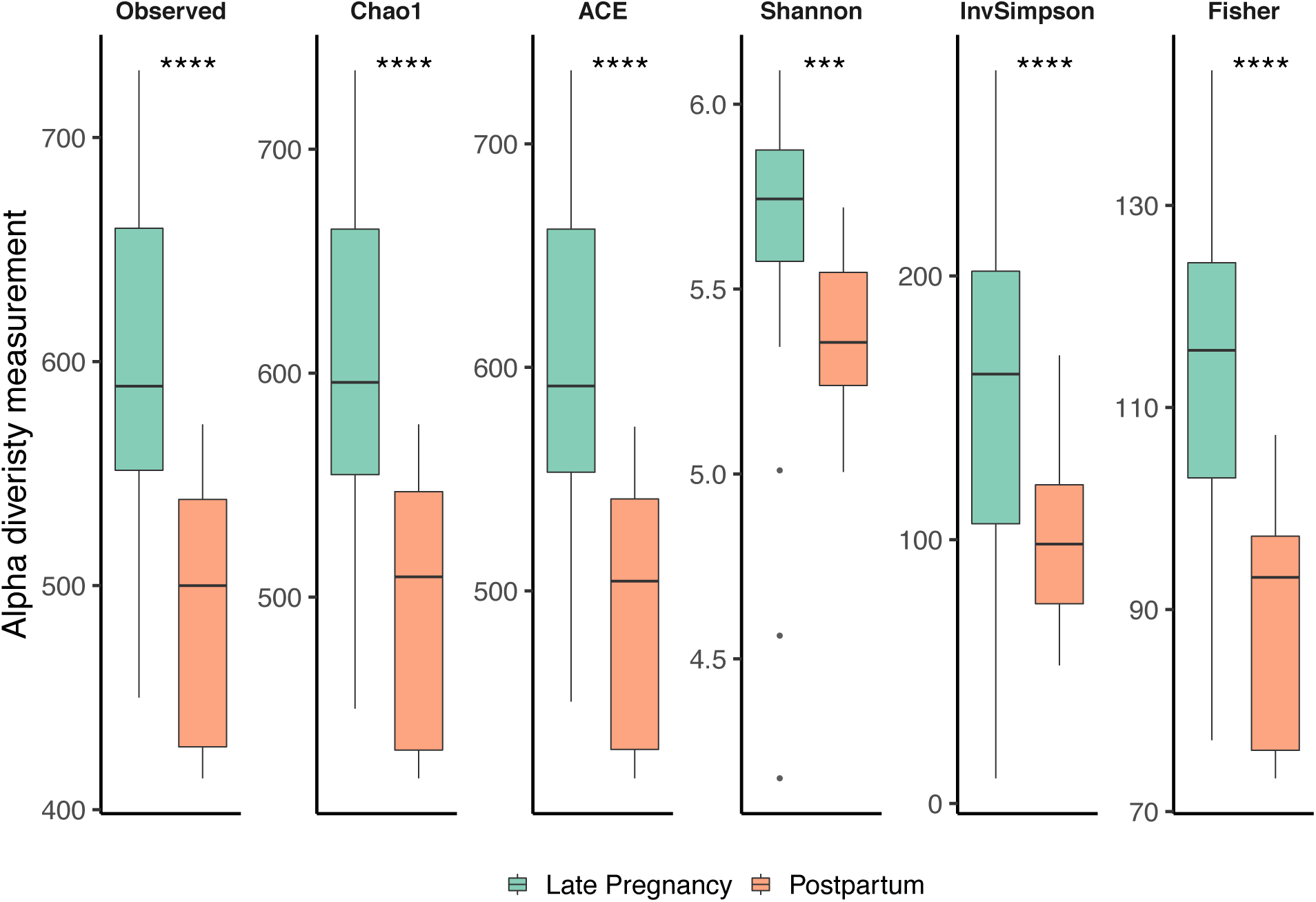
Changes in alpha diversity metrics of the maternal gut microbiome of female Tibetan antelope in different reproductive states. Statistical significance was assessed by t-tests. Note: ***: *p* < 0.001; ****: *p* < 0.0001.

### The composition of maternal gut microbiota shifts in the perinatal period

There was a clear separation in maternal gut microbial composition by female reproductive states based on the results of both hierarchical clustering (Figure 3) and PCoA analyses (Supplementary Figure 5). The differences in the microbial composition remained significant after controlling for the time of the year (*p* = 0.001). The core set of the maternal gut microbiome (including the dominant phylum Firmicutes, Bacteroidetes and Verrucomicrobia) remained stable in the transition from the late pregnancy to the postpartum period (Figure 1). However, there was significant enrichment in Actinobacteria (+1.44%; adjusted *p* < 0.05), Tenericutes (+0.42%; adjusted *p* < 0.01), Cyanobacteria (+0.48%; adjusted *p* < 0.001) and Proteobacteria (+0.15%; adjusted *p* < 0.05) during late pregnancy compared to the postpartum period (Figure 1). At the family level, *Christensenellaceae* were more abundant (+3.31%; adjusted *p* < 0.001) during late pregnancy, while *Muribaculaceae* were more abundant during the postpartum period (+1.6%; adjusted *p* < 0.001). At the genus level, *Christensenellaceae_R-7_ group* (+3.33%; adjusted *p* < 0.001), *Ruminococcaceae_UCG-010* (+2.46%; adjusted *p* < 0.01) and *Ruminococcaceae_UCG-014* (+0.95%; adjusted *p* < 0.01) were significantly enriched during the late pregnancy, whereas *Ruminococcaceae_UCG-005* (+9.04%; adjusted *p* < 0.001), *Alistipes* (1.87%; adjusted *p* < 0.001) and *Lachnospiraceae_NK4A136_ group* (+1.08%; adjusted *p* < 0.01) were significantly enriched during the postpartum period.

**Figure 5.**
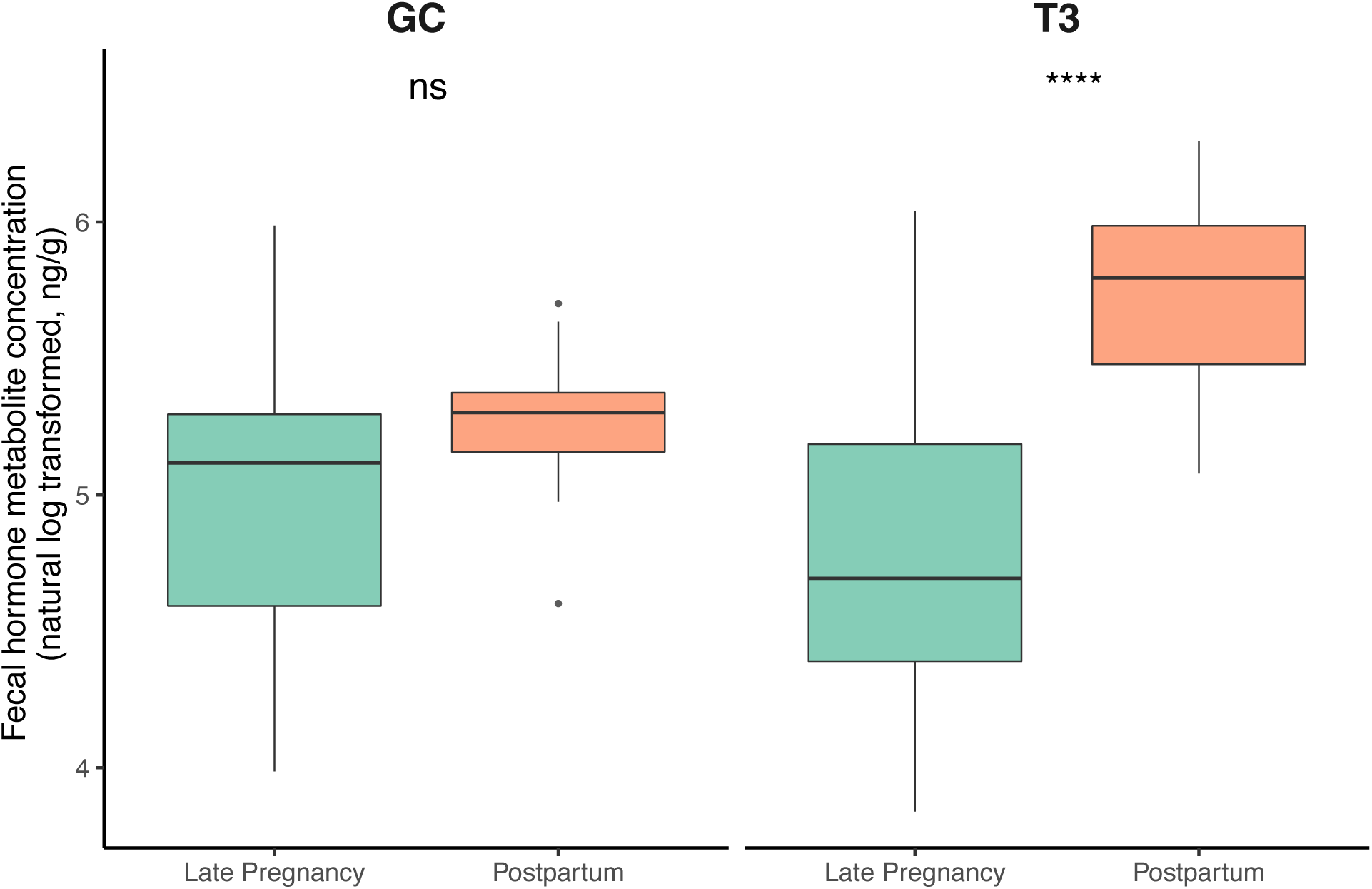
Changes in fecal GC and T3 metabolite concentrations (natural log-transformed) between reproductive states. Statistical significance was assessed by Wilcoxon signed-rank tests. Note: ns: not significant. ****: *p* < 0.0001.

### Impacts of hormonal changes on gut microbiome diversity

Overall, alpha diversity in the gut microbiota of female Tibetan antelope was significantly higher in the late pregnancy period compared to those in postpartum period across all alpha diversity indices we tested (observed number of ASVs: *p* < 0.0001; Chao1: *p* < 0.0001; ACE: *p* < 0.0001; Shannon: *p* < 0.001; InvSimpson: *p* < 0.0001; Fisher: *p* < 0.0001) (Figure 4). Fecal GC metabolite concentrations did not significantly differ between reproductive states (*p* = 0.06; Figure 5, left). However, T3 concentrations were significantly higher in the postpartum period compared to late pregnancy (*p* < 0.0001; Figure 5, right). There was a significant negative correlation between T3 and alpha diversity measurement (Observed, Chao1, ACE and Fisher) during the transition from late pregnancy to the postpartum period (*p* < 0.05; Supplementary Figure 6), however, this relationship became insignificant when only considering each reproductive state separately (Figure 6). There was no significant correlation between GC and any microbiome alpha diversity measurement (Figure 6 and Supplementary Figure 6). T3 only explained 5.70% of the total variance in microbiome alpha diversity among samples (*adj R*^*2*^ = 0.057). When we included both T3 and the reproductive states in the linear model, the fit of the model increased with *adj R*^*2*^ as 0.306.

**Figure 6.**
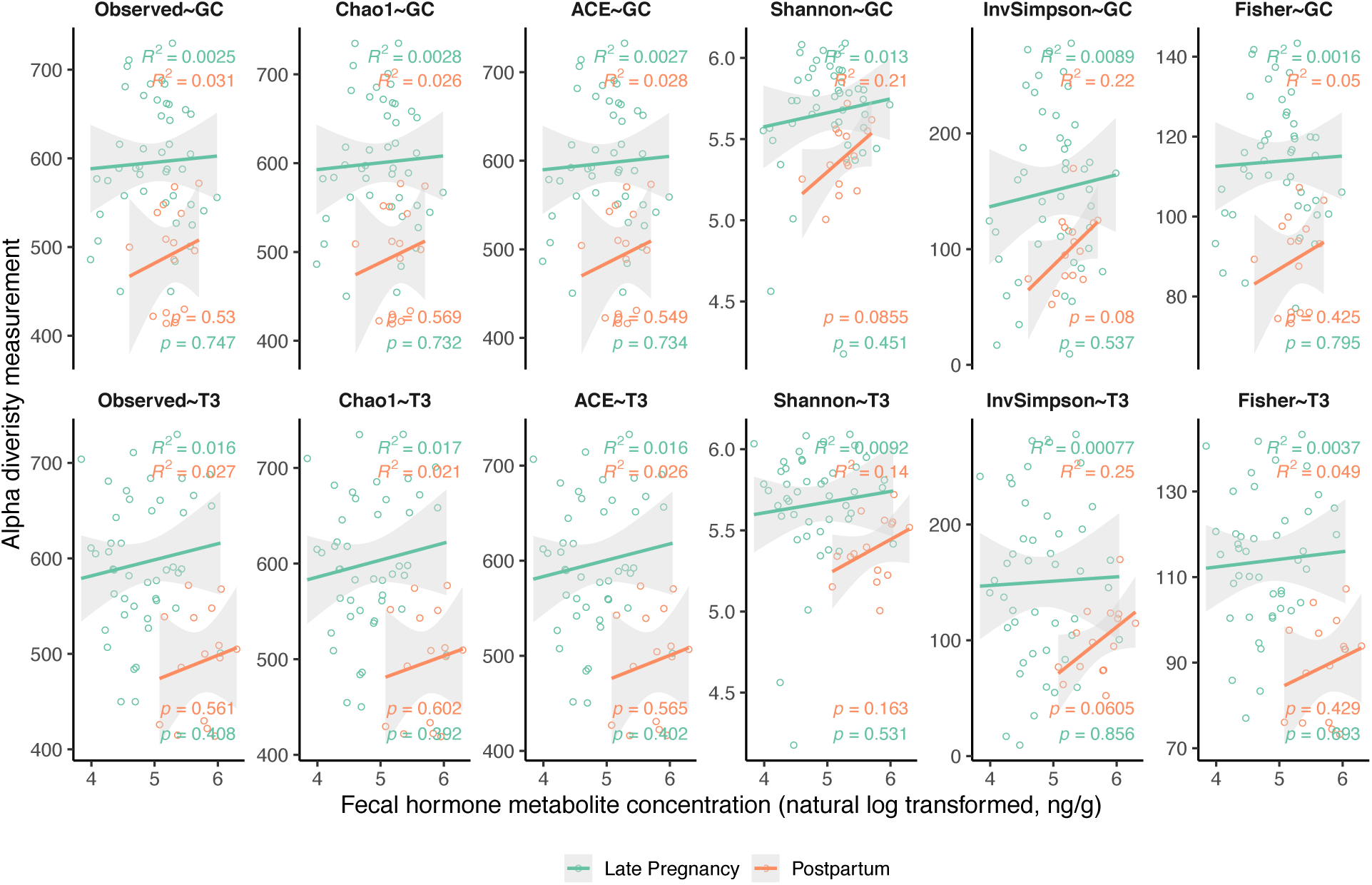
Relationships between fecal hormone metabolite concentrations (GC and T3) and microbiome alpha diversity measurements at different reproductive states.

## Discussion

In this study, we characterized the maternal gut microbiota of Tibetan antelope and revealed a significant shift in the microbiome community composition and reduced microbiome diversity in the transition from late pregnancy to the postpartum period. We also found that there was a significant negative correlation between T3 and microbiome diversity (Observed, Chao1, ACE and Fisher) during this transition, however, this relationship became insignificant when considering each reproductive state separately.

The maternal gut microbiota of Tibetan antelope is dominated by Firmicutes and Bacteroidetes (Figure 1). This finding is consistent with the microbiome composition in other mammals (Lau et al., 2018; Ley et al., 2008). In herbivores, these two dominant phyla accounted for 79% −86% of the total microbiome community (O’ Donnell et al., 2017). Firmicutes and Bacteroidetes are strict anaerobes and dominate the deeply anaerobic environments such as animal guts (Friedman et al., 2018). Facultative anaerobes, such as Proteobacteria and Actinobacteria, are typically 100-fold lower in abundance compared to strict anaerobes (Nagpal et al., 2017). Another vital role played by the symbiont gut microbiome is nutrient uptake (Krajmalnik-Brown, Ilhan, Kang, & DiBaise, 2012). Animals are capable of digesting proteins, lipids, and simple sugars through enzymatic breakdown. However, these enzymes are not able to digest the complex structural polysaccharides of plants (Dearing & Kohl, 2017). Microbes can break down these structural polysaccharides into volatile fatty acids, allowing them to be absorbed and assimilated to provide energy for the host (Dearing & Kohl, 2017). The average ratio between Firmicutes and Bacteroidetes in Tibetan antelope is 2.52:1, which corresponds to the trade-offs between carbohydrate and protein fermentation. Plant-based diets increase the relative abundance of Firmicutes that metabolize dietary plant polysaccharides while decreasing the relative abundance of Bacteroidetes that is closely associated with protein-based diets (David et al., 2013; Lima et al., 2015). Verrucomicrobia is the third most abundant phylum (2.94%) in the gut microbiome of Tibetan antelope (Figure 1) and is reportedly important for hydrolyzing multiple polysaccharides, including laminarin, xylan and chondroitin sulfate (Cardman et al., 2014). The majority of Verrucomicrobia-associated sequences were classified to the genus *Akkermansia*, which are biomarkers for a healthy mucus layer in the animal gut and bring a competitive advantage during nutrient deprivation (Belzer & de Vos, 2012). In total, these three phyla had a combined relative abundance of 97.21%. This core set of maternal gut microbiome represents a significant benefit for Tibetan antelope in terms of energy intake and nutrition acquisition. We also found the presence of Cyanobacteria in the gut microbiome community of Tibetan antelope, though its prevalence was only 0.472% (Figure 1). Cyanobacteria have only been recently recognized in the gut microbiome and are specifically common in the guts of herbivores (Di Rienzi et al., 2013). Recently, these Cyanobacteria-related gut bacteria have been assigned to a new phylum, Melainabacteria. In addition to helping with the digestion of plant fibers, Melainabacteria can also synthesize several B and K vitamins (Di Rienzi et al., 2013).

There was a significant reduction in the microbiome alpha diversity from late pregnancy to the postpartum period (Figure 4), consistent with a previous study (Crusell et al., 2018). We also found distinct microbiome profiles during late pregnancy and the postpartum period (Figure 3). This finding is not surprising, considering the postpartum period plays a vital role in “resetting” maternal changes accumulated during pregnancy (Stuebe & Rich-Edwards, 2008). Visceral fat accumulates and insulin resistance level increase throughout gestation (Lain & Catalano, 2007; Nelson, Matthews, & Poston, 2010). The accumulated fat stores are mobilized post-partum to support lactation, shifting resource allocation from energy storage to milk synthesis. Lactation results in improved insulin sensitivity, drop in inflammation and reduced adiposity. Microbiome might play a crucial role in this transition, ensuring continuous energy supply to support the growth and development of the fetus and lactation. We found an increased abundance of members of Actinobacteria and Proteobacteria during late pregnancy, as previously reported (Koren et al., 2012). Actinobacteria are positively related to the plasma glucose level (Crusell et al., 2018) and may increase insulin insensitivity during pregnancy (Koren et al., 2012). Proteobacteria are known to include multiple pathogens, leading to proinflammatory changes and altered gut microbiota in favor of dysbiosis (Mukhopadhya, Hansen, El-Omar, & Hold, 2012; Shin, Whon, & Bae, 2015). A normal healthy pregnancy is characterized with controlled mild maternal systemic inflammatory response, with increased leukocytes and intracellular reactive oxygen species (Sacks, Studena, Sargent, & Redman, 1998). The maternal proinflammatory environment promotes the contraction of the uterus, the expulsion of the baby and rejection of the placenta during the stage of late pregnancy (Mor, Cardenas, Abrahams, & Guller, 2011). But heightened maternal inflammation during pregnancy can have adverse impacts on the offspring, including increased risk of brain development problems (Rudolph et al., 2018).

*Ruminococcaceae_UCG-005* was significantly enriched during the postpartum period and they are found to be positively related to elevated concentration of acetate, butyrate and total SCFA (Gao et al., 2019). Taken collectively, these changes in microbiome composition represent reflect the metabolic and immune changes in the critical transition period from late pregnancy to the postpartum period. Unexpectedly, we found *Christensenellaceae* were more abundant during late pregnancy. Members of *Christensenellaceae* are reportedly associated with lean host phenotype (Goodrich et al., 2014), however, pregnancy is characterized with excess adiposity and weight gain. We only started to decipher the functions of the gut microbiome in the perinatal period and many unknowns remain to be explored. For example, the family *Muribaculaceae* is recently classified (Lagkouvardos et al., 2019) and we are not sure about their precise roles in the transition from late pregnancy to the postpartum period. The same applies to other taxa, such as *Christensenellaceae_R-7_ group, Ruminococcaceae_UCG-010, Ruminococcaceae_UCG-014, Alistipes*, and *Lachnospiraceae_NK4A136_ group*. Additional in-depth genetic and functional studies will be required to understand their roles in this transition period.

We found higher T3 levels in the postpartum period (Figure 5). T3 is positively correlated with metabolism, with reduced T3 slowing down the metabolism to guard against the body using up its remaining reserves, and *vice versa* (Douyon & Schteingart, 2002). Majority studies have shown similar basal metabolic rates in the lactating and pregnancy state (Butte & King, 2005) and relatively constant T3 level in this transition period (Hendrick, Altshuler, & Suri, 2011). Study in wild Amazon river dolphin showed that lactating and non-pregnant adult females had significantly higher total T3 concentrations than pregnant females, and this difference was primarily driven by the drop in the total T3 concentrations during the late pregnancy, likely due to competition for circulating iodine from the fast-growing fetus (Robeck et al., 2019). The observed increased T3 level in the postpartum period in our study might suggest an increase in energy intake and/or increase in energetic demands in migratory lactating female Tibetan antelope. In fact, their return trip takes twice as long as the trip migrating to the calving grounds (Buho et al., 2011). Unexpectedly, we did not observe a reduction in GC level in the postpartum period (Figure 5) as in humans. Maternal plasma GC level increases throughout human pregnancy, reaching a peak near term (Concannon, Butler, Hansel, Knight, & Hamilton, 1978). The GC level declines towards the pre-pregnancy level after delivery as the HPA axis gradually recovers from its activated state during pregnancy (Hendrick et al., 2011; Mastorakos & Ilias, 2003). However, the rate of HPA normalization period varies from a couple of days to a couple of months in humans (Abou-Saleh, Ghubash, Karim, Krymski, & Bhai, 1998; Glynn, Davis, & Sandman, 2013; Jung et al., 2011; Mastorakos & Ilias, 2003). Sampling time might explain the reduced GC level in the postpartum period of Chiru. It takes Tibetan antelope about 8 days to reach the calving ground from our sampling location at Wubei Bridge. The animals usually stay at the calving ground for 8-20 days, and the return trip takes about 14-16 days (Buho et al., 2011). Since we did not collect samples immediately before parturition, we may have missed the samples with the peak GC level. By the time we collected the postpartum samples, the GC level could have dropped to the degree that is comparable to the level in the pregnant samples we collected.

We did not find a significant correlation between hormonal changes (T3 or GC) and the microbiome diversity when considering each reproductive state separately (Figure 6). T3 only explained 5.70% of the total variance in microbiome alpha diversity among samples. When we included both T3 and the reproductive states in the linear model, the fit of the model increased to 30.6%. Therefore, the observed change in the microbiome diversity was mainly and significantly driven by the reproductive transition in the linear models. However, our results do not rule out the impacts of metabolic hormones on microbiome diversity, because there was a small but significant negative correlation between T3 and alpha diversity measurement (Observed, Chao1, ACE and Fisher) during the transition from late pregnancy to the postpartum period (Supplementary Figure 6). We only focused on the perinatal period, which was a relatively short period (about 45 days). The lack of correlation between T3 and microbiome diversity right before and after parturition should not be generalized to the whole gestation and lactation period. There might be other hormones worth investigating in the future studies, such as insulin, insulin-like growth factor, C-peptide, glucagon, gastrointestinal polypeptide, ghrelin, leptin, and resistin (Gomez-Arango et al., 2016). However, these hormones are peptide hormones and can only be measured with serum samples. Future studies should also look into metabolomics, as hormones might affect metabolic activities of the microbiome as well.

One assumption we made throughout this study is that fecal samples collected in May - June were collected from pregnant females (during westward migration to the calving ground), and the samples collected in July - August (during eastward migration back to the wintering ground) were from postpartum females. Since only female Tibetan antelope conduct seasonal long-distance migration (Schaller, 1998), we were confident that there were no adult male samples in our collection. We acknowledge that not all migratory females are pregnant and a proportion of migratory individuals are non-pregnant yearlings (Schaller, 1998). In the postpartum period, the ratio of adults to young females is about 2:1 based on field observations (Schaller, 1998; Xia et al., 2007). We avoided collecting fecal samples from the young and newborns by ignoring smaller sized fecal pellets. Future studies could include progesterone in the suit of fecal hormonal metabolite assays to filter out nonpregnant migratory females. However, there are no fecal endocrine measures that can identify lactating females (Hodges & Heistermann, 2011).

The gut microbiome can respond rapidly to dietary change (David et al., 2013) and thus seasonal dietary change could be a confounding factor in this study. However, the differences in the microbial composition remained significant after controlling for the time of the year (*p* = 0.001). The migration period of Tibetan antelope overlaps with the short plant-growing season on the Tibetan Plateau (Mo et al., 2018). The forage quality is relatively consistent throughout the migratory period, and only drops after September (Leslie & Schaller, 2008). We collected fecal samples at the same location when female Tibetan antelope cross the Wubei Bridge during the late pregnancy and postpartum period, so the vegetation composition in their diets was relatively consistent. Therefore, the impacts of dietary changes on the microbiome during the migration period were likely minimal compared to the shift in reproductive states.

In conclusion, we characterized the maternal gut microbiota of wild Tibetan antelope and demonstrated its shift during the transition from late pregnancy to the postpartum period. The shift appears to support energetic demands and immune system modulation of these two reproductive states. Microbiome diversity was significantly reduced in the postpartum period. Neither GC nor T3 significantly affected microbiome diversity when only considering each reproductive state separately. Further investigations on other metabolic hormones or immune system are needed to clarify underlying reasons for the microbiome community shift and reduced microbiome diversity during the perinatal period. Our study is the first case to integrate microbiome analysis into wildlife reproduction research with an emphasis on the importance of such integration on reproductive health and overall recovery success for priority species.

## Acknowledgements

We thank Rebecca Booth for the advice on hormonal assays. We thank Kekexili Natural Nature Reserve Administration for assistance in the fieldwork. We thank Richard G. Olmstead and Noah Synder-Mackler for feedback that greatly improved the manuscript. The study was funded by the China Scholarship Council (CSC) Graduate Research Fellowship, Fritz/Boeing International Research Fellowship and WRF Hall Fellowship.

## Data Accessibility

After peer review and before final publication, the raw sequence data will be deposited in NCBI under the SRA accession number #########. The following data will be deposited in Dryad: (i) filtered ASV sequences (*fastq* format); (ii) filtered ASV abundance table with taxonomic affiliations (*csv* file); (iii) sample metadata information (*csv* file).

## Author Contributions

YS, JPS, and SKW conceived the project. YS designed the experiments. YS and ZYM conducted the experiments. YS conducted the analyses and wrote the manuscript. SKW edited the manuscript. All authors declare no conflict of interest.

