## Supplementary for "Shift of maternal gut microbiome of Tibetan antelope (*Pantholops hodgsonii*) during the perinatal period"

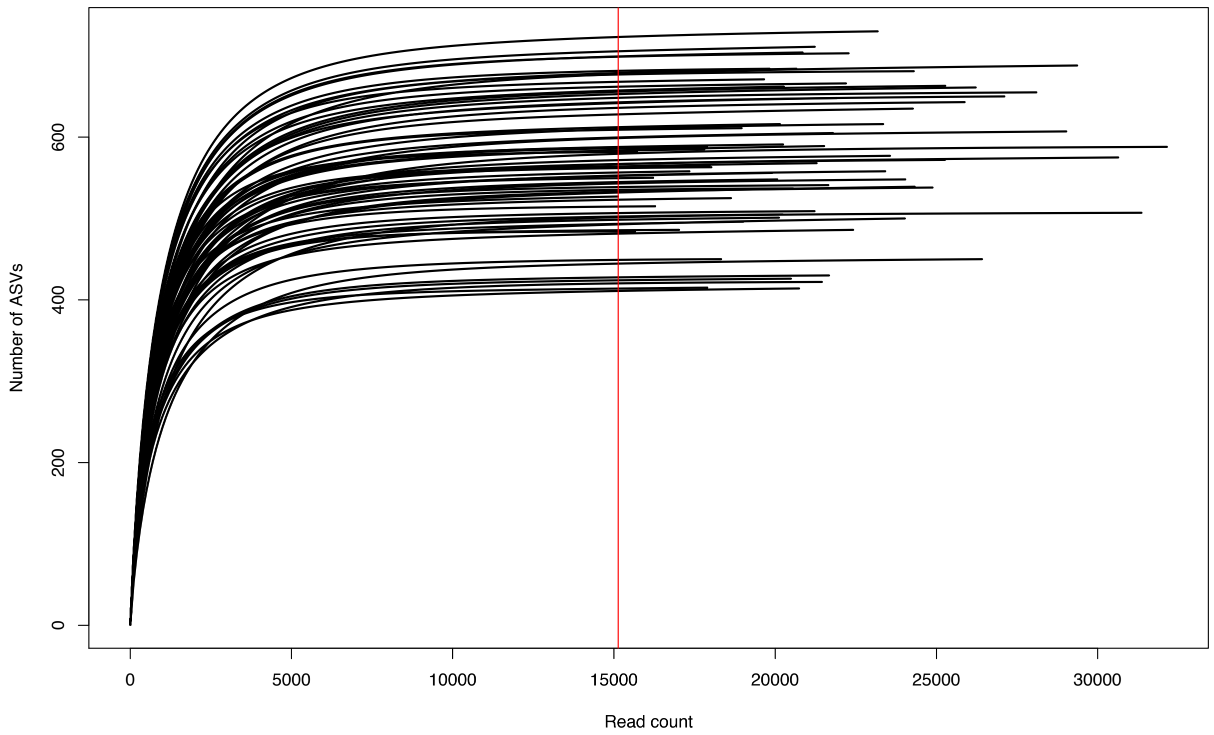


**Supplementary Figure 1** Rarefaction curves calculated for the number of amplicon sequence variants (ASVs) with increasing sequencing depth. Note: each curve represents a sample and N=65. The red vertical line indicates the minimum number of reads found in the dataset after filtering, which is 15,129 reads.

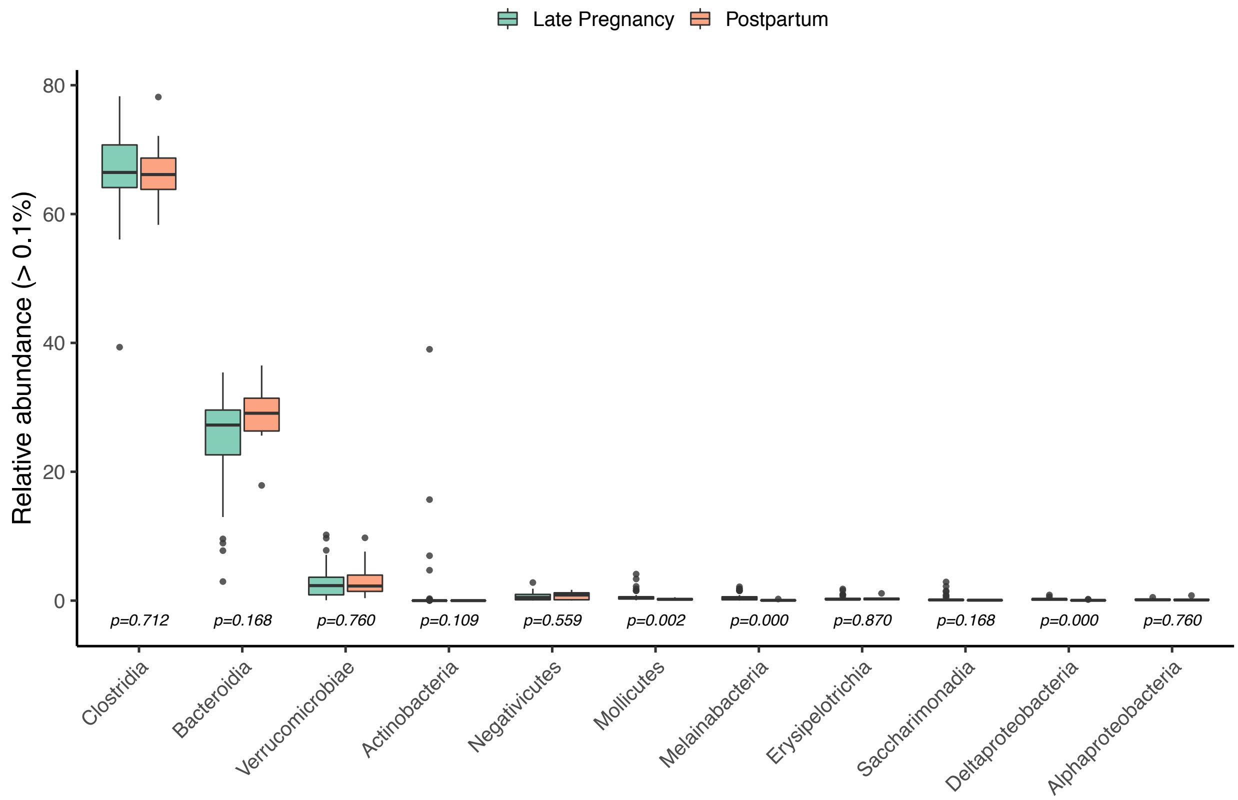


**Supplementary Figure 2** Classes found in the maternal gut microbiome of Tibetan antelope with relative abundance greater than 0.1%. Changes in the relative abundance of classes between reproductive states were analyzed through Wilcoxon signed-rank tests and $p$ values were adjusted with the Benjamini-Hochberg method to control for false discovery rate.

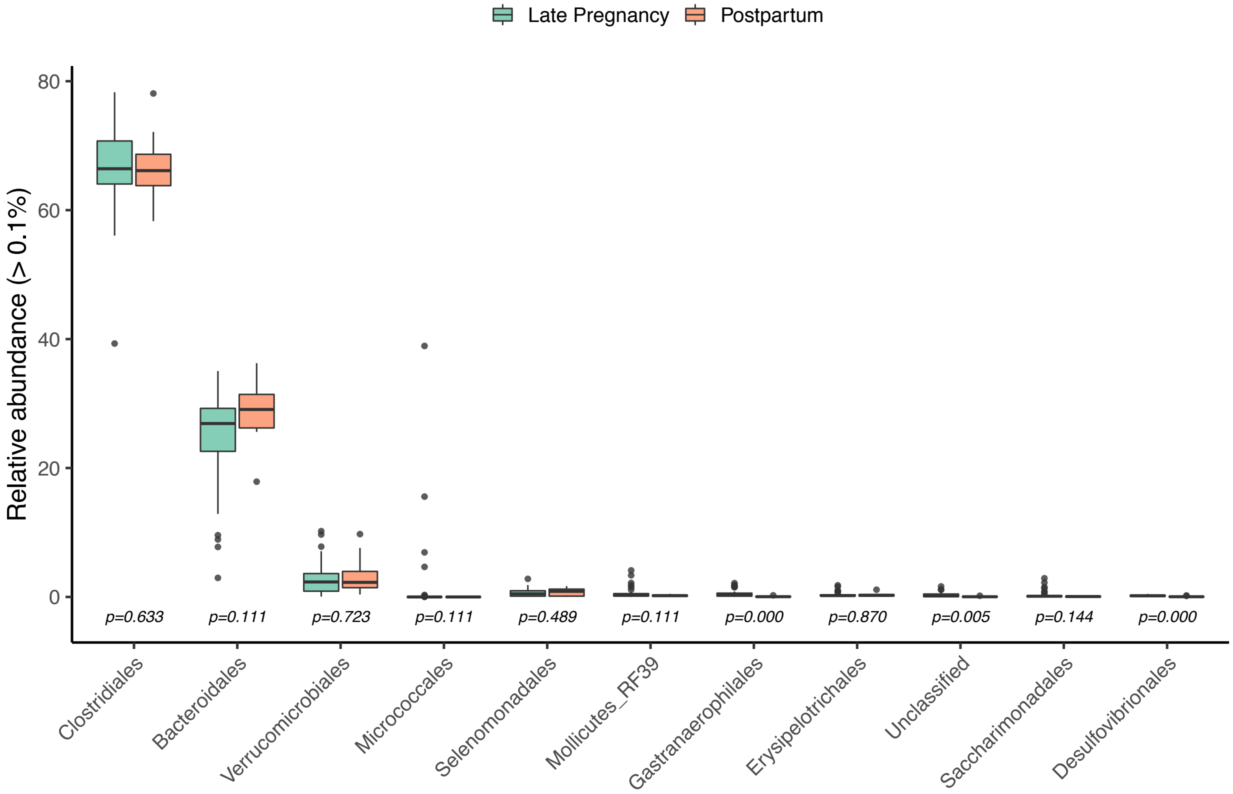


**Supplementary Figure 3** Orders found in the maternal gut microbiome of Tibetan antelope with relative abundance greater than 0.1%. Changes in the relative abundance of orders between reproductive states were analyzed through Wilcoxon signed-rank tests and *p* values were adjusted with the Benjamini-Hochberg method to control for false discovery rate.

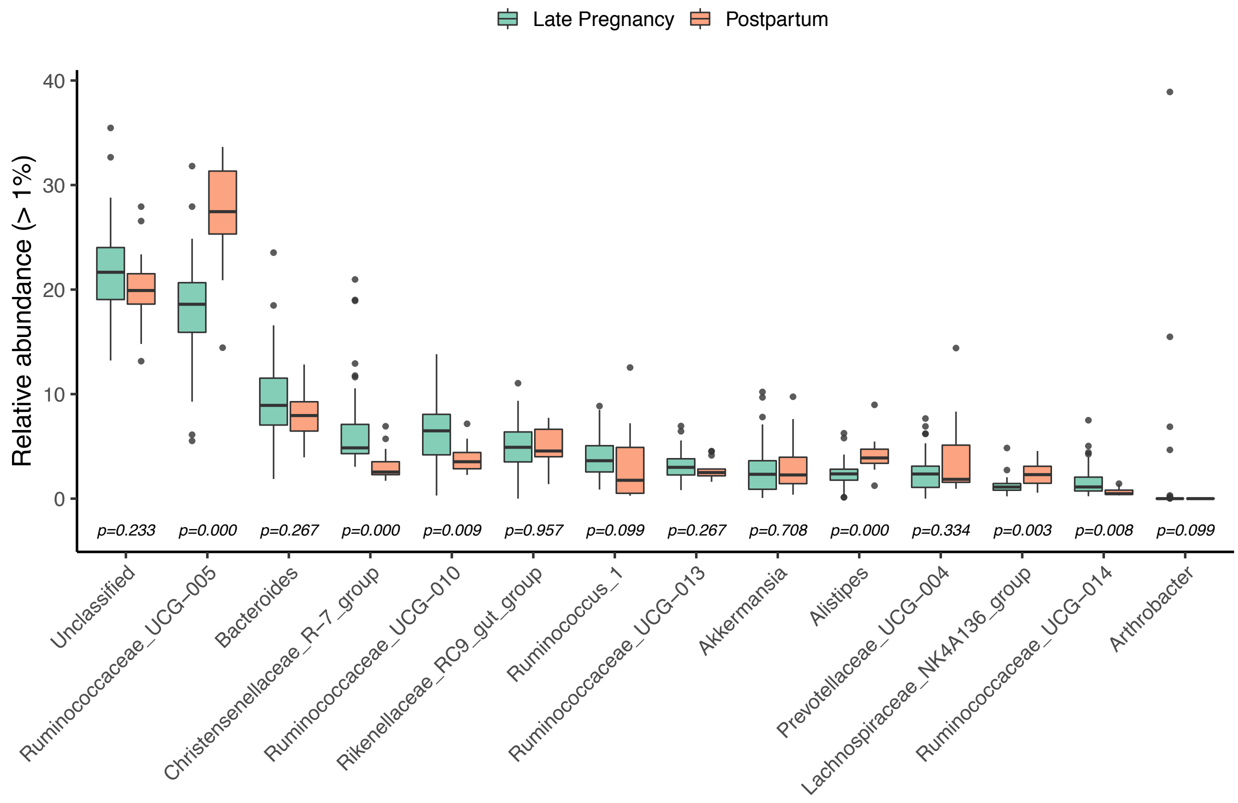


**Supplementary Figure 4** Genera found in the maternal gut microbiome of Tibetan antelope with relative abundance greater than 1%. Changes in the relative abundance of genera between reproductive states were analyzed through Wilcoxon signed-rank tests and *p* values were adjusted with the Benjamini-Hochberg method to control for false discovery rate.

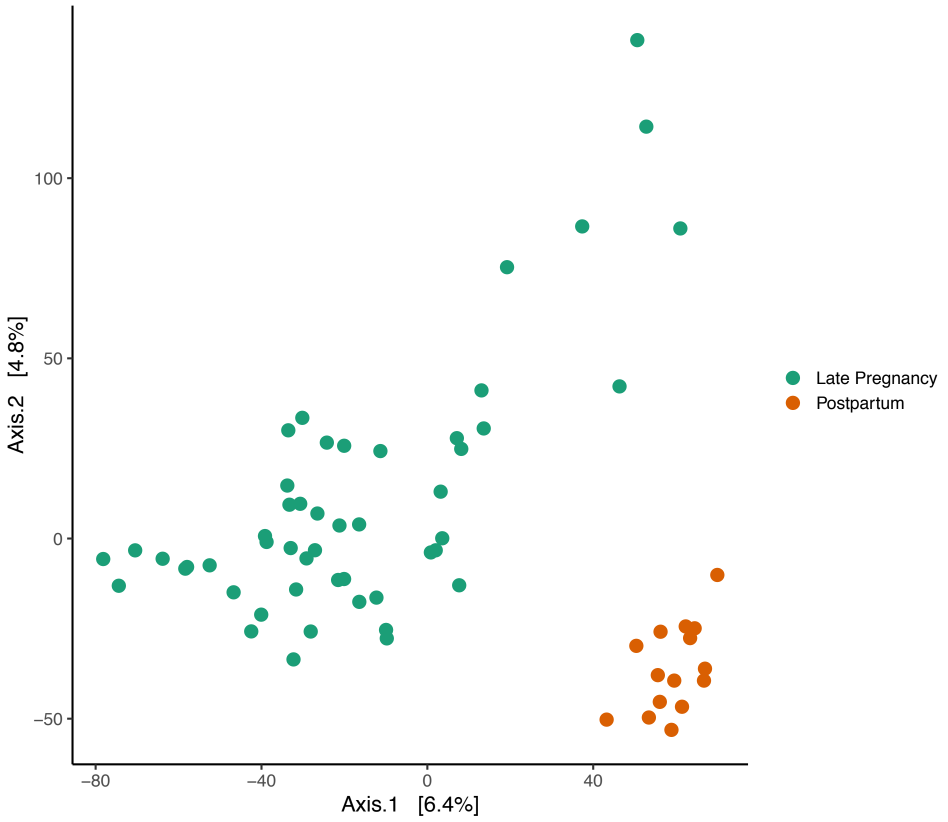


**Supplementary Figure 5** Principal coordinates analysis (PCoA) for gut microbial communities as a function of the reproductive state (N=65). The analysis was based on Euclidean distance after variance stabilizing transformation.

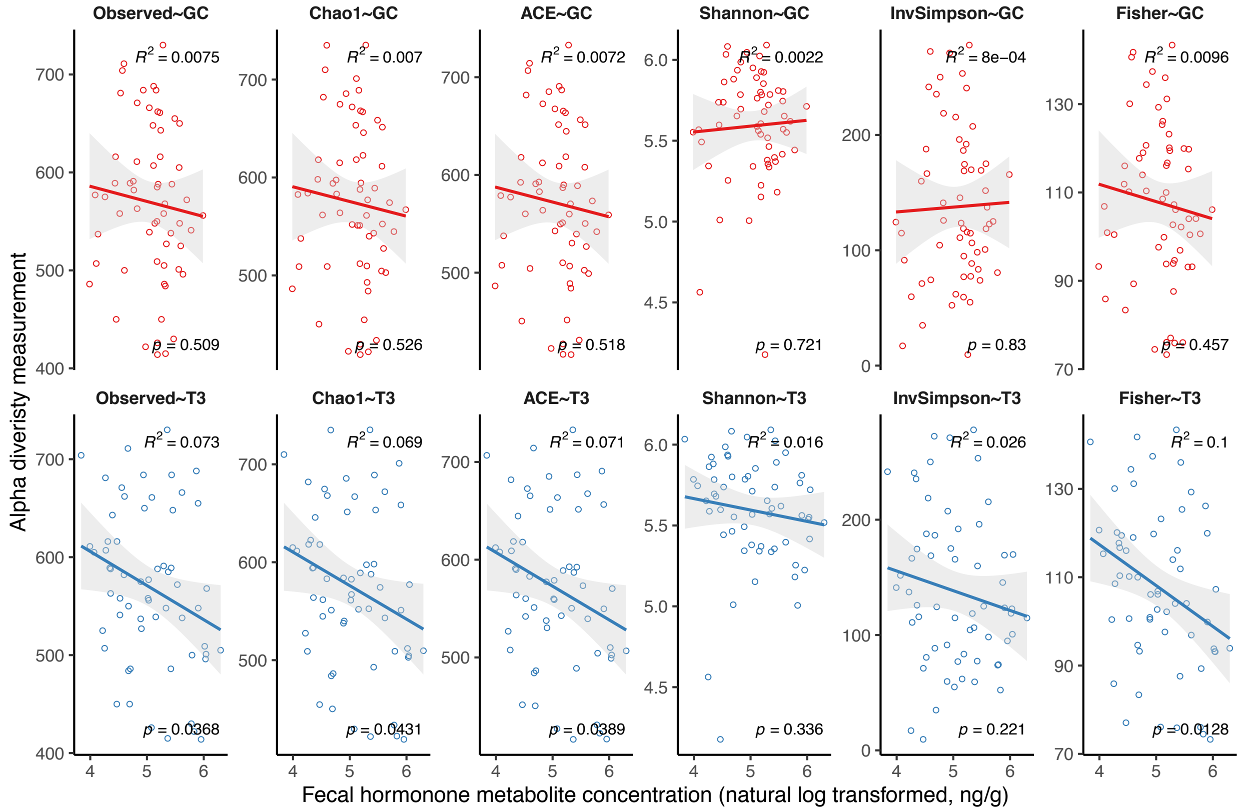


**Supplementary Figure 6** Relationships between fecal hormone metabolite concentrations (GC and T3) and microbiome alpha diversity measurements regardless of reproductive states.
